# Inflammasome-Inhibiting Nanoligomers are Neuroprotective Against Space-Induced Pathology in Healthy and Diseased 3D Human Motor and Pre-Frontal Cortex Brain Organoids

**DOI:** 10.1101/2024.03.19.585802

**Authors:** Sadhana Sharma, Vincenzo S. Gilberto, Jon Rask, Anushree Chatterjee, Prashant Nagpal

**Affiliations:** Sachi Bio, 685 S Arthur Avenue, Colorado Technology Center, Louisville, CO 80027, USA; NASA Ames Research Center, Moffett Field, California, CA 94035, USA

**Keywords:** Space-induced Neurodegeneration, Nanoligomer, Brain Organoids, Alzheimer’s Disease (AD), Amyotrophic Lateral Sclerosis (ALS), 3D Human Organoids, microgravity, space biology, International Space Station

## Abstract

Microgravity and space environment has been linked to deficits in neuromuscular and cognitive capabilities, hypothesized to occur due to accelerated aging and neurodegeneration in space. While the specific mechanisms are still being investigated, spaceflight-associated neuropathology is an important health risk to space astronauts and tourists, and is being actively investigated for the development of appropriate countermeasures. However, such space-induced neuropathology offers an opportunity for accelerated screening of therapeutic targets and lead molecules for treating neurodegenerative diseases. Here we show, a proof-of-concept high-throughput target screening (on Earth), target validation, and mitigation of microgravity-induced neuropathology using our Nanoligomer™ platform, onboard the 43-day SpaceX CRS-29 mission to the International Space Station (ISS). First, comparing 3D healthy and diseased pre-frontal cortex (PFC, for cognition) and motor neuron (MN, for neuromuscular function) organoids, we assessed space-induced pathology using biomarkers relevant to Alzheimer’s Disease (AD), Frontotemporal Dementia (FTD), and Amyotrophic Lateral Sclerosis (ALS). Both healthy and diseased PFC and MN organoids showed significantly enhanced neurodegeneration in space, as measured through relevant disease biomarkers, when compared to their respective Earth controls. Second, we tested the top two lead molecules, NI112 which targeted NF-κB, and NI113 that targeted IL-6. We observed that these Nanoligomers significantly mitigate the AD, FTD, and ALS relevant biomarkers like amyloid beta-42 (Aβ42), phosphorylated Tau (pTau), Kallikrein (KLK-6), Tar DNA-binding protein 43 (TDP-43), and others. Moreover, the 43-day Nanoligomer treatment of these brain organoids did not appear to cause any observable toxicity or safety issues in the target organoid tissue, suggesting good tolerability for these molecules in the brain at physiologically relevant doses. Together, these results show significant potential for both the development and translation of NI112 and NI113 molecules as potential neuroprotective countermeasures for safer space travel, and demonstrate the usefulness of the space environment for rapid, high-throughput screening of targets and lead molecules for clinical translation. We assert that the use of microgravity in drug development and screening may ultimately benefit millions of patients suffering from debilitating neurodegenerative diseases on Earth.

## INTRODUCTION

### The Spaceflight Environment Causes Neuromuscular and Cognitive Deficits

Several studies on astronauts have reported a decrease in neuromuscular function, cognition, poor spatial orientation and coordination, as a result of prolonged space travel.^1–5^ To quantify these deficits, NASA’s neurocognition assessment tool “*Cognition*” is used by astronauts to evaluate any changes in cognitive abilities using a battery of tests.^6^ As reported from the *NASA Twin Study*, cognitive assessment through this tool showed clear deficits for the subject after 340 days in space, compared to his preflight baseline as well as his twin brother (as Earth control).^7^ Microstructural changes in gray and white matter, vestibular and sensorimotor alterations,^1–5^ low-dose galactic cosmic radiation-induced protein misfolding, accelerated neuropathogenesis,^1,8^ and decrements in motor skills and cognitive performance were all observed.^9,10^ Assessment of these deficits from a molecular biology and pharmacology perspective is essential for the development of appropriate countermeasures for safer space travel.

Interestingly, we discovered that the conditions of the spaceflight environment also offer a unique opportunity where space offers conditions that mimic the acceleration of disease progression for screening and rapid drug discovery for multiple neurodegenerative diseases. There have been several catalyst events like FDA-incentivization for removing the animal-based testing,^11^ as well as several high-profile failures of long-awaited neurotherapeutic drugs like Aduhelm, highlighting the lack of reliability of target ID and early translation in animal-based models. Because microgravity accelerates aging and neurodegeneration in human organoids, a large opportunity for drug discovery in the space economy has emerged.^12^ However, to date, there have been very few studies validating the disease-based biomarkers that can quantify the need and specific pharmacological intervention needed as an effective countermeasure. Such studies could also provide a benchmark on what neurodegenerative disease on Earth can benefit from neurodegenerative acceleration. One possible example is Amyotrophic Lateral Sclerosis (ALS) since both microgravity exposure and ALS are associated with significant motor abnormalities, exhibiting similarities in the degradation of the spinal cord, genetic markers, and pathogenesis.^13–15^ Similarly, microgravity and space cause a cognition deficit due to accelerated neurodegeneration centered around a combination of oxidative stress and mitochondrial dysfunction leading to increased inflammation,^16^ oxidative species-induced protein misfolding, and plaque buildup,^8,16^ microgravity-induced neuropathy^1^ mimicking Alzheimer’s Disease (AD).

### Neuroinflammation Targets for Multiple Neurodegenerative Diseases

The NASA twin study documented the spaceflight-induced low-grade upregulation in inflammatory cytokines, shortened telomeres, and other attributes of accelerated aging,^7^ resembling characteristics of “inflammaging”.^17,18^ Role of activated microglia and astrocytes has already been well-documented in the initiation and progression of several neurodegenerative diseases.^19–22^ Various parts of the immune system are involved in disease initiation and progression of neuropathology, ranging from activation of immune cells CD4+ T cells (involving NF-κB transcriptional factor), to the release of pro-inflammatory cytokines such as tumor necrosis factor-alpha (TNF-α), IL-1β, and Interleukin 6 (IL-6), that contribute to neuronal tissue damage. This abnormal immune activation also involves activation of the inflammasome pathways (e.g., NOD-like receptor family pyrin domain containing 3 (NLRP3), NLRP1), has been actively investigated and associated with a wide range of inflammatory, autoimmune, cardiometabolic, and neurodegenerative diseases.^21,23–33^ As a result, microgravity and space environment appears to stress cells in ways that cause them to rapidly develop disease pathology that mimics debilitating motor and cognitive neurodegenerative diseases like AD and ALS.^1,8,13–15^ Due to the large number of potential targets involved, a high-throughput and targeted screening method is required to identify the best therapeutic target.

### Nanoligomer-based High-Throughput Screening is a Safe and Targeted Approach

Nanoligomer™ therapeutic platform generates safe, targeted,^34^ and highly-specific (K_D_ 3.37 nM^35,36^) RNA-targeting molecules for facile delivery to targeted organs and both up-and down-regulation of desired protein through translational and transcriptional regulation.^37–42^ Nanoligomers utilize peptide nucleic acids (PNAs) as the sequence-specific binding component that demonstrate strong hybridization and specificity to their RNA (or DNA) sequence targets,^43^ and exhibit no known enzymatic cleavage, leading to increased stability in human blood serum and mammalian cellular extracts.^44^ Given the low K_D_,^35,36,43^ high binding specificity, minimal off-targeting,^35,36^ lack of any immunogenic response or accumulation in first pass organs,^34^ absence of any observable histological damage to organs even for long (>15-20 weeks) treatments,^34–36,40^ and facile delivery to the brain to counter neuroinflammation,^34,37–42^ we have demonstrated that Nanoligomers are a safe and targeted approach to screen for therapeutic targets in organoids and rodents for multiple neurodegenerative diseases aboard the International Space Station (ISS).

## RESULTS AND DISCUSSION

### High-Throughput *in-vitro* Screening in Donor-Derived Primary Human Astrocytes

Neuroinflammation mitigation consists of many potential targets. Therefore, due to limitations on the number of samples flying in space aboard the ISS, targeted screening and down selection was done on Earth using reactive and activated astrocytes (or type A1) astrocytes.^45^ Activated microglia that are responsible for neurodegenerative diseases^19–22^ have been shown to increase reactive A1 astrocytes via the release of pro-inflammatory cytokines. Using an established stimulation cocktail of IL-1α, TNF α, and C1q (complement protein), primary human astrocytes were activated to mimic the neurotoxic inflammation and induce an A1 phenotype.^45^ For high-throughput screening, Nanoligomers targeting different neuroinflammation genes (NLRP3, Nuclear factor kappa-light-chain-enhancer of activated B cells (NF-κB), tumor necrosis factor (TNF) receptor 1 (TNFR1), NLRP1, TNF, and IL-6) were used for the treatment of A1 astrocytes (see Methods), and the effect of the treatment was monitored by measuring the expression of 65-proinflammatory cytokines and chemokines (**Fig. 1A-D**). As a preliminary screen, IL-1α was used to rank the impact of Nanoligomer targets in this *in vitro* neuroinflammation model (**Fig. 1A**, gene expression data using Nanoligomer treatments shown previously^37,38^). By using Principal Component Analysis (PCA) to present the combined outcome from these 65 inflammation biomarkers, we used the primary component (PC1) representing 72.87% variance in the data, to rank these targets alongside IL-1α (**Fig. 1B**).^37,38^ Using these key parameters, and impact on 65 cytokine and chemokines (**Fig. 1C, D**), NF-κB and IL-6 were the top targets identified in the screen. The choice of NF-κB target was significant, especially because despite its importance for pharmacological intervention, it has been considered an “undruggable” target.^46,47^

**Fig. 1.**
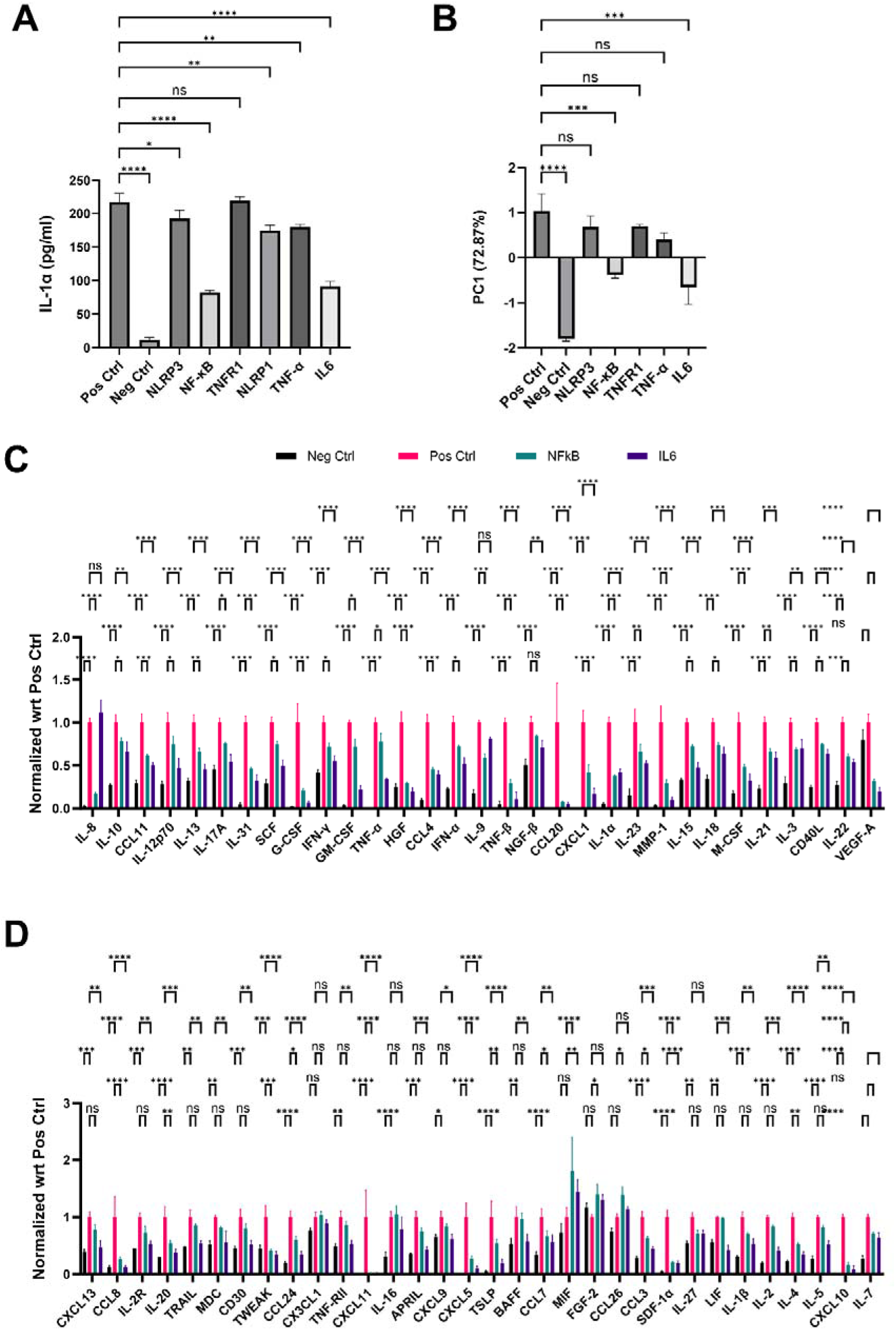
Neurotherapeutic drug-discovery and high-throughput screening for down selection of target molecules. **A**. IL-1α protein expression for untreated and treated activated A1 astrocytes, along with resting state/non-activated astrocytes as negative control. Different Nanoligomer treatments targeting separate neuroinflammatory pathways were used to screen and rank selected inflammasome-targeting Nanoligomers. **B**. Multiplexed 65 human cytokine and chemokine expression was analyzed using PCA. Primary PC1 (72.87 % variability) was used for ranking the Nanoligomers. **C**. and **D.** 65-cytokines and chemokine shown for resting state astrocytes (negative control, Neg Ctrl), activated but sham-treated astrocytes (positive control, Pos Ctrl), and top 2 ranked Nanoliogmers NF-κB and IL-6 treated astrocytes, normalized by positive control (Pos Ctrl is 1). \**P* < 0.05, \*\**P* < 0.01, and \*\*\**P* < 0.001, \*\*\*\**P* < 0.0001, Mean ± SEM, significance based on one-way ANOVA. *n* = 3 for each group.

### Assessment of Space-Induced Neuropathology in Healthy and Diseased 3D Motor Neuron (MN) and Pre-Frontal Cortex (PFC) Brain Organoids

We used the 43-day SpaceX Commercial Resupply Service (CRS)-29 mission to the ISS as a means to investigate the impact of space-induced neuropathology in healthy and diseased brain organoids (**Fig. 2A**). Due to the preponderance of observations in motor and cognitive deficits observed in astronauts, we prepared two types of healthy and diseased brain organoids using established literature techniques to culture 3D organoids.^48,49^ We used a healthy 90:10 neuron: astrocyte ratio to create 3D healthy MN organoids and PFC organoids (with 70:30 distribution of glutamatergic: GABAergic neurons). To fully capture the impact of space environment on potential cognition deterioration, we also cultured diseased PFC organoids (DPFC), using APOE4/E4 mutation (two alleles of APO4) which is the highest genetic risk factor and most prevalent in AD patients (APOE4 carriers account for 65–80% of all AD cases), and has been shown to induce detrimental effects via amyloid Aβ plaque, neurotoxic fragments, stimulating Tau phosphorylation, and impaired mitochondrial function.^50–56^ In parallel, we also used TDP-43 overexpression (TDP-43 knock-in) in neuronal cells to create diseased MN organoids (or DMN), to induce frontotemporal dementia (FTD) and ALS-mimicking pathology in 3D MN organoids. TDP-43 has been shown to be a key risk and pathological factor for both FTD and ALS, as well as other neurodegenerative diseases.^57,58^

**Fig. 2.**
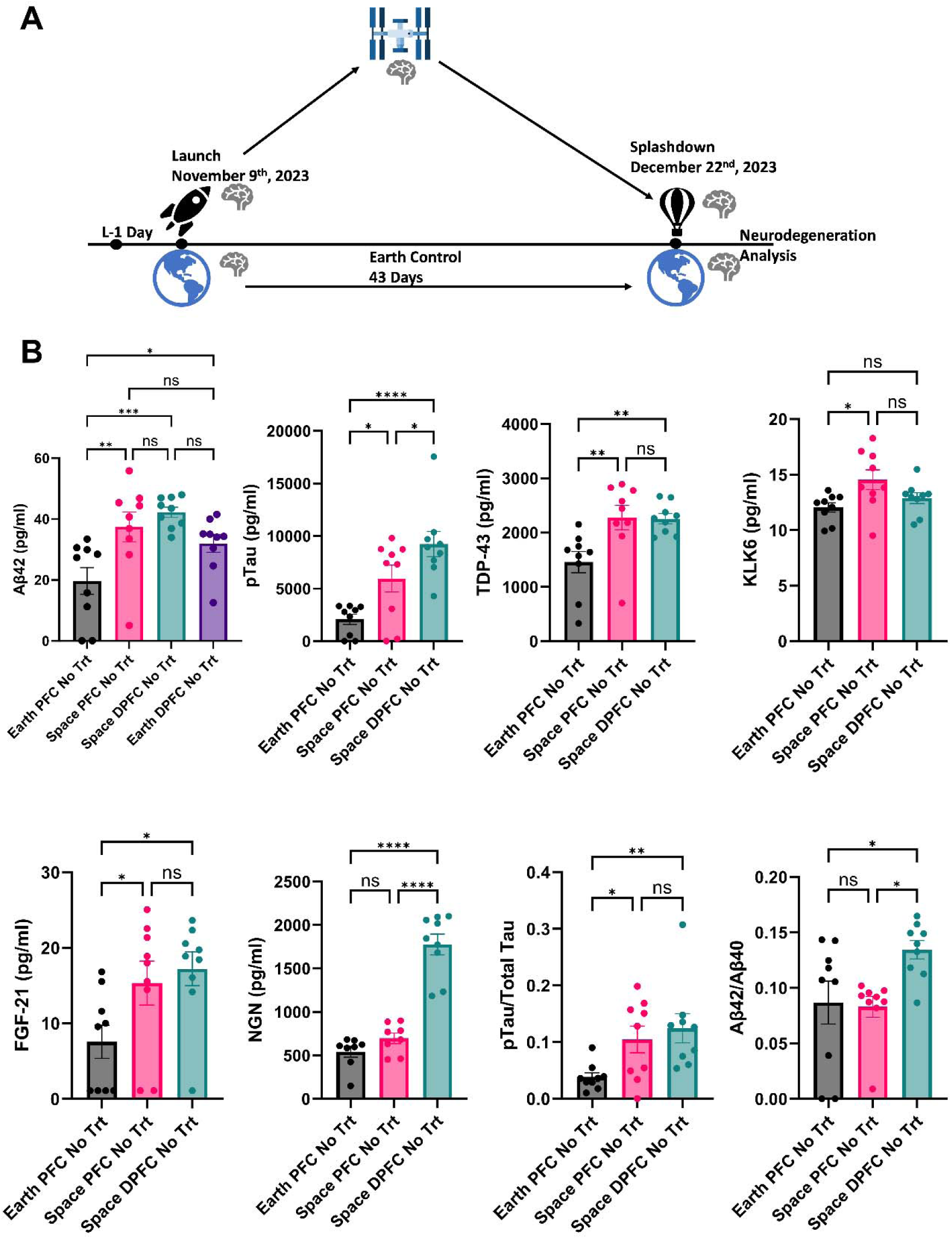
Comparison between healthy PFC and diseased (DPFC) 3D human organoids in Space (at ISS) and on Earth (ground control) to evaluate space-induced neuropathology. **A**. Schematic showing ISS-based drug discovery on CRS-29, a 43-day spaceflight mission. 3D human brain organoids were used to evaluate the effect of space-induced pathology on neurodegeneration, and the potential reversal of disease biomarkers through screened Nanoligomer molecules. **B.** Using the following neurodegenerative biomarkers, healthy PFC and diseased DPFC organoids were compared between the ISS (Space) and ground controls (Earth): Aβ42, pTau, TDP-43, KLK6, FGF-21, NGN, pTau/Total Tau, and Aβ42/ Aβ40. All biomarkers show significant upregulation of neurodegeneration, as evidence of space-induced neuropathology. Each dot represents an individual organoid measurement. \**P* < 0.05, \*\**P* < 0.01, and \*\*\**P* < 0.001, \*\*\*\**P* < 0.0001, Mean ± SEM, significance based on one-way ANOVA. *n* = 9 for each group.

To capture the potential induction of disease pathology for a multitude of neurodegenerative diseases, we measured nine key biomarkers in organoids flown aboard the ISS and compared them with corresponding Earth (ground) controls. They included TDP-43 expression, a neurodegenerative biomarker related to diseases that cause both motor and cognitive deficits (e.g. ALS, AD),^57–60^ Amyloid Beta 42 (Aβ42) the constituent found in brain plaques that is directly linked to AD disease pathology of cognitive decline and dementia,^61^ and phosphorylated tau (pTau) which has been linked to the formation of neurofibrillary tangles (NFTs)^62^. To expand the repertoire to other diseases and additional neuropathologies, we also measured concentrations of Kallikrein-6 (KLK6), Fibroblast Growth Factor 21 (FGF-21), Aβ42, Aβ40, pTau, total Tau (Total Tau), Neurogranin (NGN), and neural cell adhesion molecule (NCAM-1) using multiplexed ELISA (**Fig. 2B**). For example, FGF21 has been shown to ameliorate neurodegeneration in rodent and cellular models of AD.^63^ KLK6 is shown to be directly linked to axonal and neuronal degeneration and motor function loss,^64^ as well as other neuropathologies such as neuroinflammation and Tau phosphorylation leading to identification in other CNS diseases too.^65^ NGN that is a post-synaptic protein has been used as a clinical biomarker and observed to significantly increase with aging-related cognitive decline as well as in AD patients.^66,67^ Similarly, NCAM-1 protein which has been associated with synaptic plasticity has now been shown to have a role in neuropathological conditions such as AD, MS, Schizophrenia, and others.^68^ Besides cognition, many of these biomarkers have also been implicated and used in the detection of neuromuscular diseases like ALS, MS, and others. Typically, while elevated amyloid plaques Aβ42^69^ and pTau (pT181)^70^ levels are nominally associated with plaques and neurofibrillary tangles as AD pathologies, evidence of increased levels of these proteins have also been documented in neuromuscular diseases like ALS, MS, and others.^69,70^ Similarly, pTau and total Tau,^71^ TDP-43,^58,72^ NCAM-1,^68^ and NGN^67^ have well-documented roles as broader neurodegeneration biomarkers. Therefore, we selected these clinically accepted diagnostic biomarkers to determine if space-induced neuropathology or accelerated neurodegeneration occurred.

We hypothesized that spaceflight would induce neuropathology in organoids similar to the pathology observed in neurological diseases on Earth. We tested this hypothesis by comparing untreated ground controls (Earth PFC No Trt, Earth DPFC No Trt in **Fig. 2B**; Earth MN No Trt, and Earth DMN No Trt in **Fig. 3**), to the corresponding ISS spaceflight samples (Space PFC No Trt, Space DPFC No Trt in **Fig. 2B**; and Space DMN No Trt in **Fig. 3**). We note that due to space limitation on samples that could fly aboard ISS on the CRS-29 space mission, untreated MN controls could not be added (Space MN No Trt), and we compared diseased MN (DMN) on ground and ISS. Analysis of Aβ42 (**Fig. 2B**) expression noted a statistically significant increase in healthy PFC on ISS after a 43-day ISS mission (comparing untreated Earth PFC and Space PFC, increase ∼2-fold, **Fig. 2B**), but only a nominal increase in diseased PFC which was not statistically significant (comparing untreated Earth DPFC and Space DPFC). Comparing other key AD biomarkers like TDP-43, KLK6, and pTau/total Tau, we saw the same effect where healthy PFC levels in space are statistically significantly higher than their healthy PFC ground controls, but similar to DPFC in space (**Fig. 2B**). However, pTau, NGN, and Aβ42/ Aβ40 levels were elevated when comparing PFC and DPFC in space, indicating more longer-term initiation of phosphorylated tau and resultingly normalized Aβ42 levels could possibly result in across the board higher biomarker pathology for even longer missions. However, significant increases in observed neurodegeneration and disease biomarkers for both healthy and diseased PFC organoids showed clear evidence of space-induced neuropathology.

**Fig. 3.**
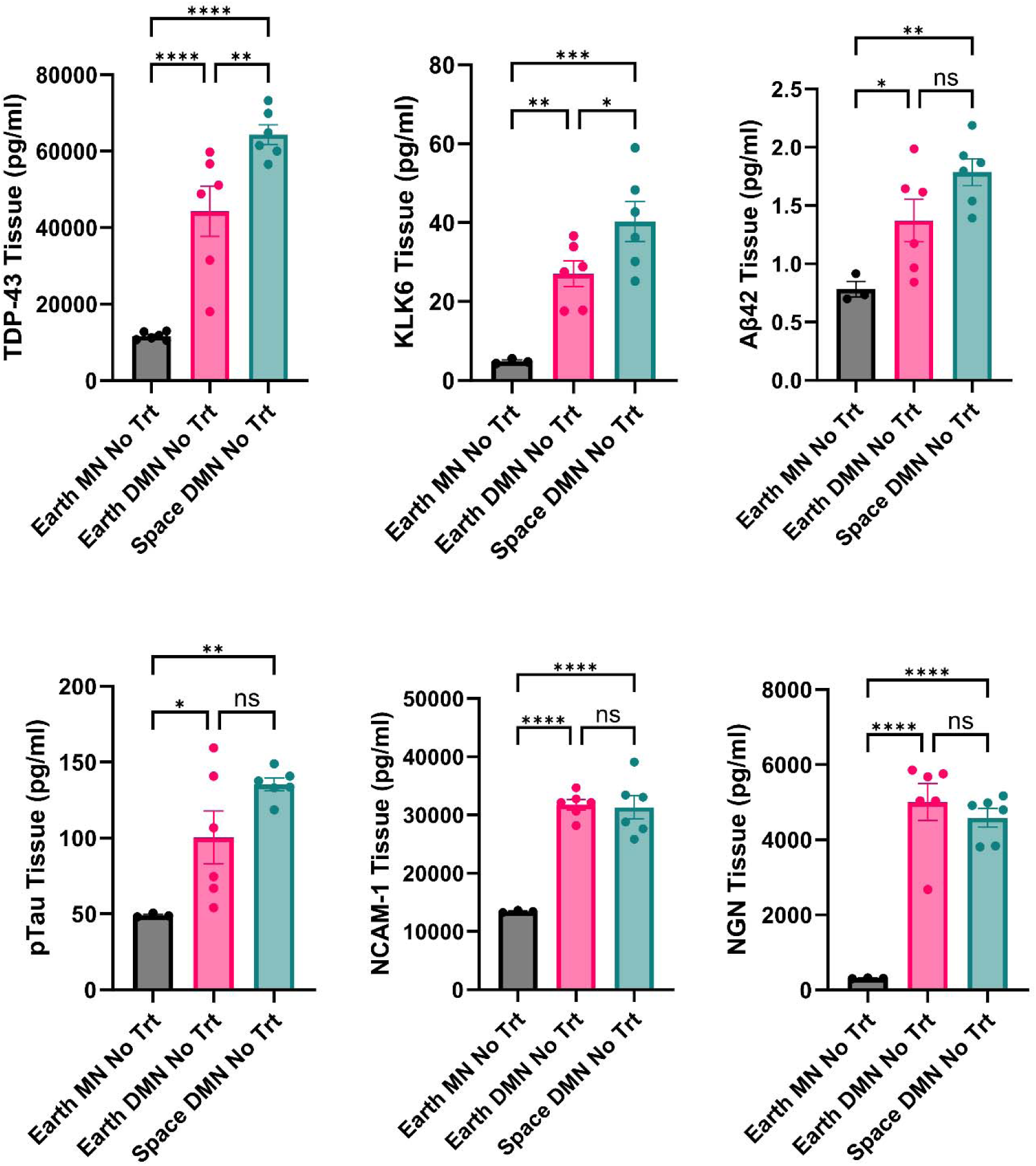
Comparison between healthy MN and diseased (DMN) 3D human organoids in Space and on Earth to evaluate the potential impact on neuromuscular disease pathology. Using the following neurodegenerative biomarkers, healthy MN and diseased DMN organoids were compared between Space and Earth: TDP-43, KLK6, Aβ42, pTau, NCAM-1, and NGN. All biomarkers show significant upregulation of neurodegeneration mimicking multiple neuromuscular diseases, as evidence of space-induced neuropathology. Each dot represents combined tissue lysed from multiple organoids (with an identical number of lysed cells). \**P* < 0.05, \*\**P* < 0.01, and \*\*\**P* < 0.001, \*\*\*\**P* < 0.0001, Mean ± SEM, significance based on one-way ANOVA. *n* = 3-6 for each group.

To assess the same hypothesis for motor neurons, we compared diseased MN (DMN) on Earth to those flown in Space, using healthy Earth MN as a benchmark. In two key biomarkers most relevant for ALS and MS, we saw a statistically significant increase of TDP-43 and KLK6 in Space DMN samples as compared to Earth controls (**Fig. 3**). While Aβ42 and pTau also showed some increase, the statistical significance was not observed, whereas NCAM-1 and NGN showed nearly similar results in diseased organoids (**Fig. 3**). Collectively, our PFC and MN results reveal that the spaceflight environment aboard the ISS induces an increase in motor function-related disease biomarkers in healthy and diseased organoids.

### Nanoligomer Treatments Ameliorate Space-Induced Neuropathology in Healthy and Diseased PFC Brain Organoids

Next, we tested the hypothesis that our previously identified top Nanoligomer targets (NF-κB and IL-6) and their respective molecules (NI112 and NI113) can be neurotherapeutic, and counter the pathology induced in space. Analysis of the AD-relevant disease biomarkers we measured (Aβ42, pTau, TDP-43, NGN, Aβ42/ Aβ40, and pTau/Total Tau), NI112 (target NF-κB) revealed that treatment with Nanoligomers in space was able to reduce pathology in diseased DPFC and restore baseline biomarker expression back to values seen in healthy PFC on Earth (**Fig. 4**). We discovered that the treatment not only countered the space effect but also reversed the disease pathology induced in PFC. Further, while NI113 (target IL-6) treatment was also able to reverse pTau, TDP-43, NGN, and pTau/Total Tau expression, it showed slightly higher levels of Aβ42, indicating strong target engagement and reduction of AD pathology, but perhaps not at the same level as NI112 treatment (**Fig. 4**). On assessing these biomarkers using healthy PFC organoids in space, we observed a reversal of pTau, NGN, and pTau/total Tau neurodegenerative biomarkers by both treatments to baseline healthy Earth PFC levels (**Fig. 5**). KLK6 levels were slightly elevated, however NI113 treatment did not produce a statistically significant effect.

**Fig. 4.**
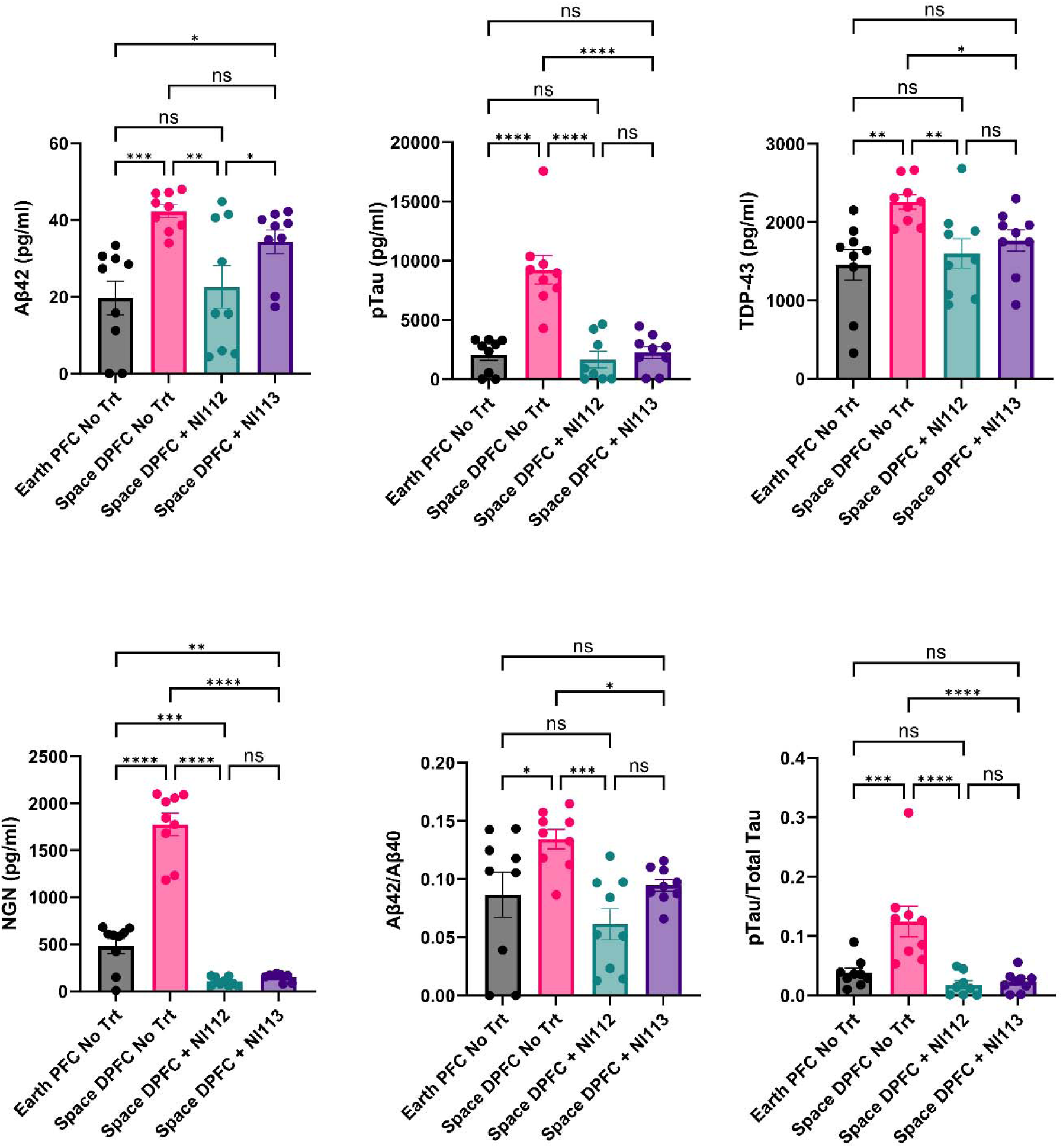
Effect of lead Nanoligomer molecules (NI112 or NI113) on diseased PFC organoids in Space, compared to healthy PFC organoids on Earth as a benchmark. Diseased PFC (DPFC) organoids in space were treated with the top two Nanoligomer lead molecules downregulating NF-κB (NI112) and IL-6(NI113), and assessed using different neurodegenerative biomarkers (Aβ42, pTau, TDP-43, NGN, Aβ42/ Aβ40, and pTau/Total Tau). Healthy PFC organoids on Earth are shown as a benchmark. All biomarkers show significant downregulation of neurodegeneration, as evidence of treatment efficacy. Each dot represents an individual organoid measurement. \**P* < 0.05, \*\**P* < 0.01, and \*\*\**P* < 0.001, \*\*\*\**P* < 0.0001, Mean ± SEM, significance based on one-way ANOVA. *n* = 9 for each group.

**Fig. 5.**
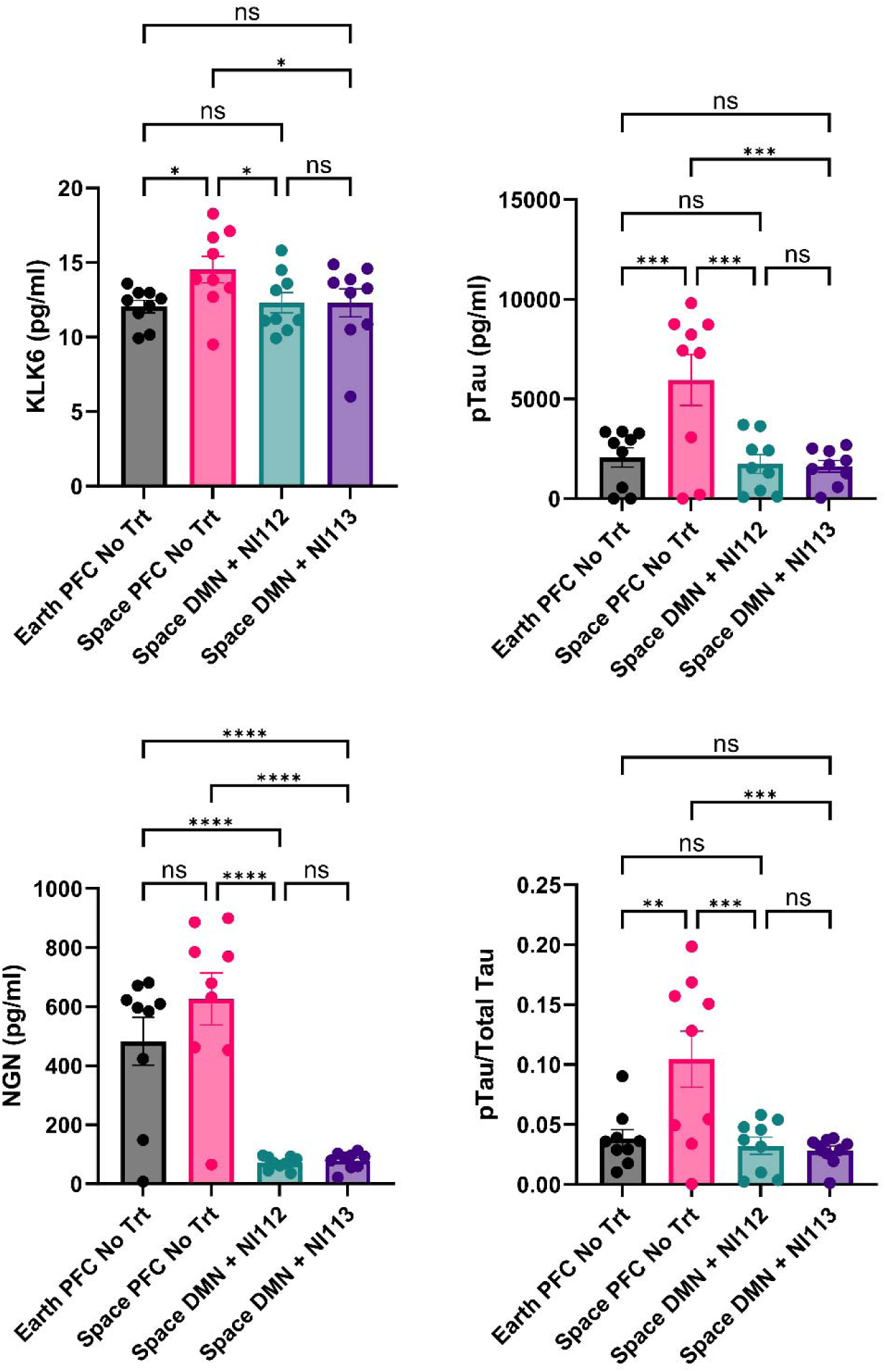
Effect of lead Nanoligomer molecules (NI112 or NI113) on PFC organoids in Space, compared to healthy PFC organoids on Earth as a benchmark. PFC organoids in space were treated with the top two Nanoligomer lead molecules downregulating NF-κB (NI112) and IL-6 (NI113), and assessed using different neurodegenerative biomarkers (KLK6, pTau, NGN, and pTau/Total Tau). Healthy PFC organoids on Earth are shown as a benchmark. All biomarkers show significant downregulation of neurodegeneration on Nanoligomer treatment. Each dot represents an individual organoid measurement. \**P* < 0.05, \*\**P* < 0.01, and \*\*\**P* < 0.001, \*\*\*\**P* < 0.0001, Mean ± SEM, significance based on one-way ANOVA. *n* = 9 for each group.

### Space-Induced Pathology is Reversed in Diseased MN Organoids with Nanoligomer Treatments

We utilized both organoid tissues as well as supernatants to measure neurodegenerative biomarkers in 3D diseased MN organoids (DMN) in Space (ISS) samples. Treatment with NI112 Nanoligomer (to target NF-κB) appeared to be significantly more effective in reducing neurodegeneration biomarkers in diseased MN in space, resulting in baseline levels similar to healthy MN on Earth (**Fig. 6**). The only exception was TDP-43 levels in Space DMN, which were still higher than Earth healthy MN controls, presumably due to strong TDP-43 expression induction genetically. IL-6 treatment (NI113 Nanoligomer) was slightly less effective and while it reduced most motor function-related biomarkers, the values were still higher compared to NI112 treatment and healthy Earth controls. We observed very similar trends in supernatant measurements where NI112 treatment seemed much more effective than NI113, whereas both Nanoliogmers significantly addressed and mitigated both space-induced pathology as well as disease induction genetically, as measured through these protein expression datasets (**Fig. 7**).

**Fig. 6.**
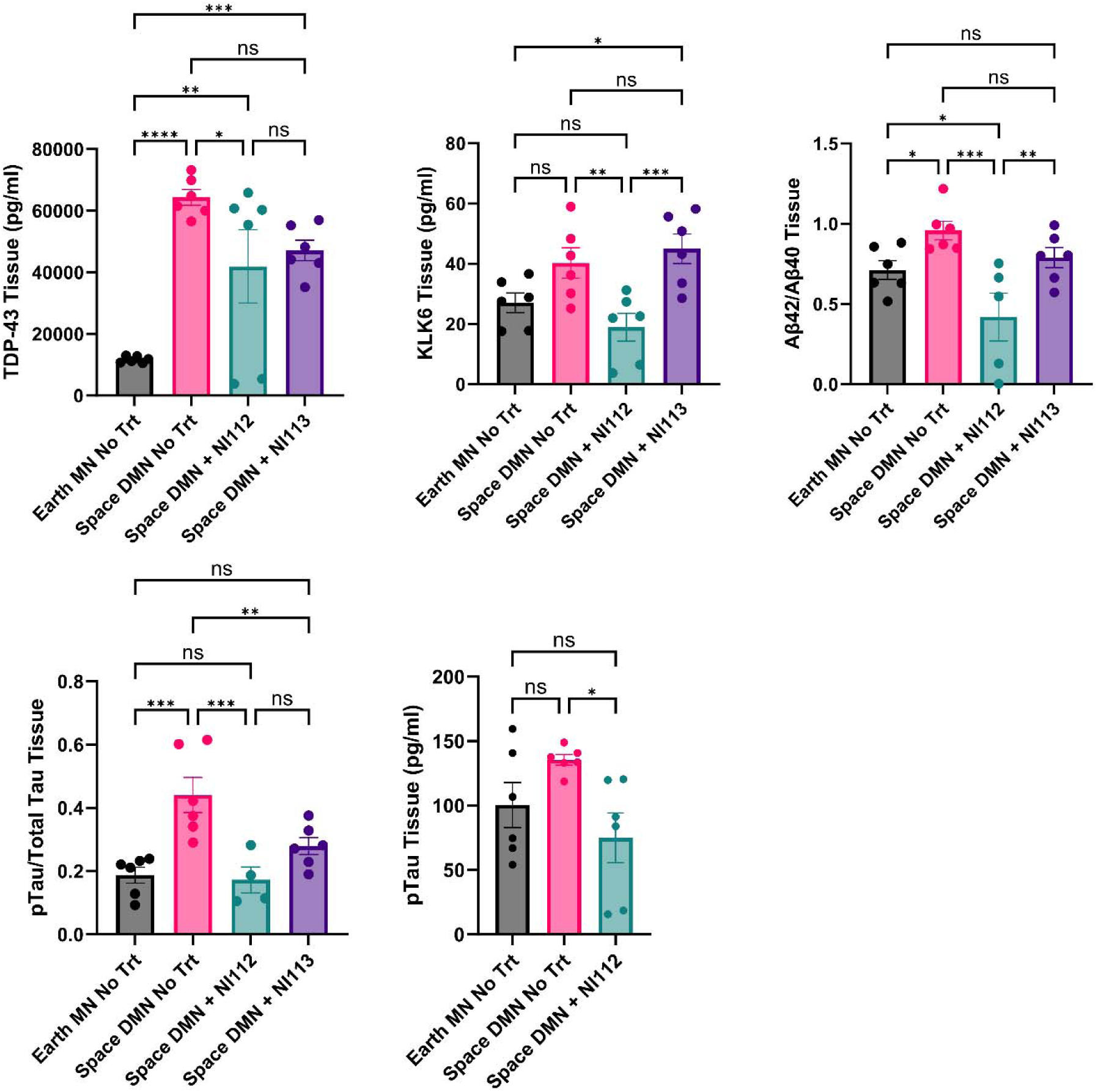
Effect of lead target (NI112 or NI113) downregulating Nanoligomer molecules on diseased MN (DMN) 3D organoids in Space, compared to healthy MN organoids on Earth as a benchmark. Tissues from diseased MN organoids were compared with (DMN + NI112, and DMN + NI113) and without (DMN No Trt) Nanoligomer treatment in space. Healthy MN organoids as ground controls (Earth) were used to benchmark treatment efficacy. All biomarkers show significant downregulation of neurodegeneration as an effect of Nanoligomer treatments. Each dot represents combined tissue lysed from multiple organoids (with an identical number of lysed cells). \**P* < 0.05, \*\**P* < 0.01, and \*\*\**P* < 0.001, \*\*\*\**P* < 0.0001, Mean ± SEM, significance based on one-way ANOVA. *n* = 6 for each group.

**Fig. 7.**
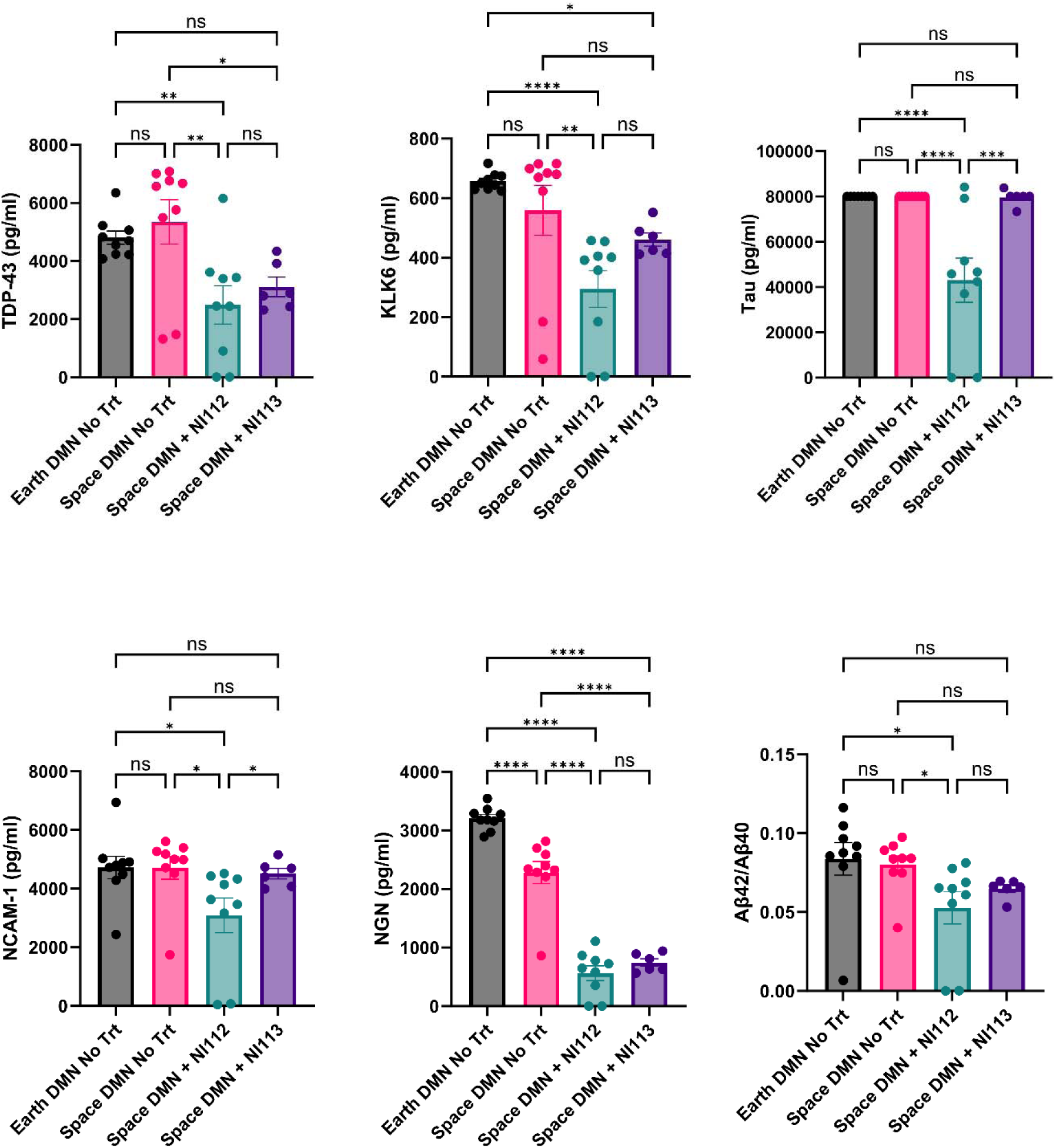
Effect of lead target (NI112 or NI113) downregulating Nanoligomer molecules on diseased MN (DMN) 3D organoids in Space, compared to healthy MN organoids on Earth as a benchmark. Diseased MN organoid supernatants with DMN + NI112, and DMN + NI113 treatments, were compared to sham-treated (DMN No Trt) organoid supernatant in space. Healthy MN organoids as ground controls (Earth) were used to benchmark treatment efficacy. All biomarkers show significant downregulation of neurodegeneration as an effect of Nanoligomer treatments. Each dot represents an organoid. \**P* < 0.05, \*\**P* < 0.01, and \*\*\**P* < 0.001, \*\*\*\**P* < 0.0001, Mean ± SEM, significance based on one-way ANOVA. *n* = 6-9 for each group.

## CONCLUSIONS

In conclusion, we used healthy and diseased PFC and MN organoids and nine disease biomarkers to assess the impact of spaceflight on potential neuropathology that is known to be related to cognition and motor function. Further, we conducted a high-throughput screen of neuroinflammation targets and identified the top 2 targets (NF-κB and IL-6) and respective downregulating Nanoligomer molecules NI112 and NI113. Both molecules showed significant downregulation of most neurodegenerative biomarkers in space. In particular, NF-κB treatment was able to reduce both space-induced and genetically induced disease-specific pathology. These results are significant since both targets do not directly address any of the biomarkers being measured, and their downregulation significantly ameliorates a wide range of disease biomarkers, indicating a significant or pivotal role neuroinflammation and inflammasome activation play in neurodegenerative disease activation. These results are in line with other observations from the NASA Twin study where a significant acceleration of aging and upregulation of inflammatory cytokines was observed, all mimicking inflammaging. This data shows that the identified lead molecules shown here can be both useful neurological countermeasures in future spaceflight experiments, as well as strong candidates for further testing in neurodegenerative diseases like AD, FTD, ALS, and progressive MS. The results also show a rapid acceleration of biomarkers in 3D human brain organoids closely mimicking these disease states and lay a foundation for further exploration and use of microgravity to advance rapid and high-throughput drug discovery in space.

## MATERIALS AND METHODS

### Nanoligomer design and synthesis

Nanoligomers were specifically designed and synthesized (Sachi Bio) following established methods detailed in published references.^35–40^ The Nanoligomers are composed of 17-base-long antisense peptide nucleic acid (PNA)^37–39,73^ conjugated to a gold nanoparticle.^37–39^ Various PNA sequences (provided in previous publications^35–40)^ were screened for their solubility, self-complementing sequences, and potential off-target interactions. The PNA portion of the Nanoligomer was synthesized on a Vantage peptide synthesizer (AAPPTec, LLC) employing solid-phase Fmoc chemistry. Fmoc-PNA monomers, wherein A, C, and G monomers were shielded with Bhoc groups, were obtained from PolyOrg Inc. Post-synthesis, the peptides were attached to gold nanoparticles and subsequently purified using size-exclusion filtration. The conjugation process and the concentration of the refined solution were monitored by measuring absorbance at 260 nm (for PNA detection) and 400 nm (for nanoparticle quantification).

### Culturing and treating Astrocytes with Nanoligomers

Primary human astrocytes (Lonza) were cultured in a complete astrocyte growth medium (ScienCell) at 37 °C and 5% CO_2,_ and passaged at ∼80-90% confluency. Cytokine (TNF-α, IL-1α, C1q-Sigma-Aldrich) stock solutions (1000X) were prepared in molecular biology grade water, aliquoted, and stored at -80°C. The effective concentrations of the various cytokines in the culture medium were: TNF-α: 30 ng/mL, IL-1α: 3 ng/mL, and C1q: 400 ng/mL.^45^ For Nanoligomer treatment, cells were seeded in a 96-well plate at a seeding density of 10,000 cells/ cm^2^ and cultured until 80% confluency. Astrocytes were then treated with the Nanoligomers for 24 hours, and a cytokine cocktail (TNF-α, IL-1α, C1q, media for negative control) for another 24 hours after which media supernatants were collected for analysis of secreted protein/cytokine expression using multiplexed ELISA.

### Culturing and treating 3D MN and PFC organoids with Nanoligomers

We used two different types of organoids: 1) MN (representing Motor cortex) and PFC (representing prefrontal cortex) area of the brain. For both MN and PFC organoids, healthy (MN, PFC) as well as diseased (DMN and DPFC) models were tested. Organoids were prepared and cultured using a slight modification of the published protocols and methods.^48,49,74^ Differentiated iCell Motor Neurons (Fujifilm Cellular Dynamics) were thawed and seeded together 90:10 neuron: astrocyte ratio in ultralow attachment (Sbio) plates and cultures for 3 weeks using complete Motor media, until the neurospheroids and organoids had matured. Typical PFC organoids/neurospheroids used 90:10 Neuron: Astrocyte ratio with 70:30 distribution of glutamatergic: GABAergic neurons.^48^ Both the diseased (DMN and DPFC), as well as healthy MN and PFC models consisted of a 90:10 Neuron: Astrocytes ratio. For the first week of growth, MN organoids were fed using Complete Motor Media with a concentration of 5uM DAPT (Cayman Chemical). Post 7 days of seeding, they were fed Complete Motor Media without DAPT. Following the 3-week maturation period, we used 200 µM Nanoligomer stock and replaced 5% of BrainPhys media with stock (10 µM final concentration). Untreated negative controls used stock media. Both Earth and Space organoids were transferred to 1mL cryovials (Nunc) using 200 µL wide bore tips (Rainin).

### Loading and Splash Down of Space Samples

Space samples were shipped from Sachi Bio, Louisville, Colorado to Kennedy Space Center. Samples were placed in a MicroQ iQ2 unit at 37°C (MicroQ Technologies). To maintain humidity Kim wipes dampened with Mili-Q water were placed in the unit along with samples. Upon arrival at the Kennedy Space Center, samples were placed in an incubator at 37°C and 5.0% CO_2_ before loading on SpaceX CRS-29. Samples were in space for 43 days. Following splashdown, Space cryovials were kept at 4°C until shipment. Corresponding Earth cryovials samples were also transferred to 4°C at the same time. Samples were then processed for supernatant collection and organoid lysis.

### Organoid lysis and protein extraction

Three organoids were combined in a 1.5 mL Eppendorf tube and washed with DPBS (Gibco). Cells were first dissociated by adding TrypLE (Gibco) and incubated at 37°C 5.0% CO2. After initial cell dissociation spheroids were neutralized and then washed. Following washing, tubes were aspirated and Bio-Plex Cell-Lysis Kit (Bio-Rad) was added and spheroids were then sonicated. Following the user’s instructions for lysis protocol, neurospheroids were then centrifuged at high speed and the supernatant was collected in a separate 200 uL PCR tube (USA Scientific).

### Single and Multiplexed ELISAs

Organoid culture supernatants and lysates were analyzed for secreted proteins using single- and multiplexed Human ProcartaPlex™ Panel (Thermofisher Scientific), following previously documented procedures.^37,38^ The plates were read using the Luminex MAGPIX xMAP instrument and xPONENT® software. Standards were prepared at 1:4 dilutions (8 standards), alongside background and controls. Subsequently, concentrations of samples were determined from a standard curve using the Five Parameter Logistic (5PL) curve fit/quantification method.

### Statistics

The figure legends specify the statistical tests conducted, the number of experiments, and the reported p-values. Microsoft Excel and GraphPad Prism 10.1.0 were employed for data analysis and data presentation.

## ASSOCIATED CONTENT

### Author Contributions

P.N. and A.C. conceived the idea and designed the experiments. P.N. synthesized the Nanoligomers. S.S., V.S.G., and A.C. cultured the organoids. A.C., S.S., V.S.G., and P.N. conducted ELISA sample preparation and measurements. P.N. wrote the manuscript with input from all the authors. All authors read the manuscript and provided input.

## ACKNOWLEDGMENTS

Authors acknowledge financial support from NASA SBIR Awards 80NSSC23CA171 and 80NSSC22PPB175, and implementation support from Space Tango.

## TOC GRAPHIC

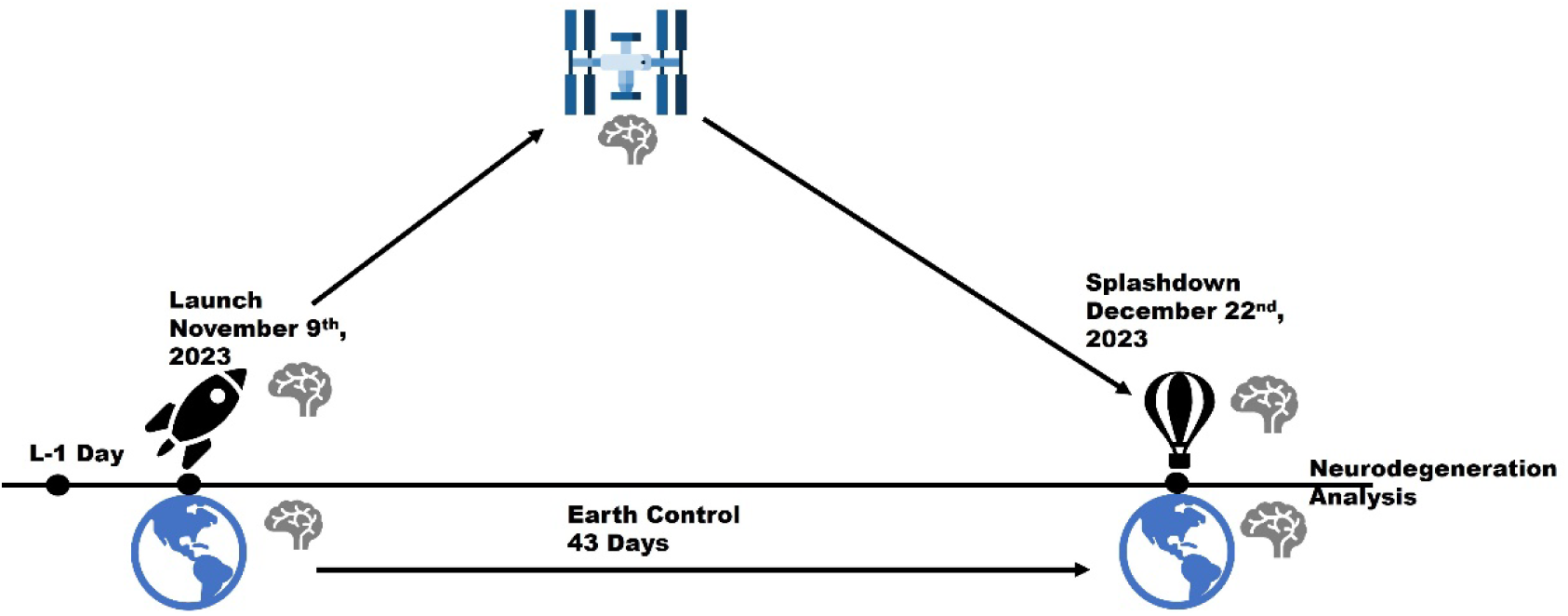

